# DBpred: A deep learning method for the prediction of DNA interacting residues in protein sequences

**DOI:** 10.1101/2021.08.05.455224

**Authors:** Sumeet Patiyal, Anjali Dhall, Gajendra P. S. Raghava

**Author notes:** Corresponding Author Prof. Gajendra P. S. Raghava, Head and Professor, Department of Computational Biology, Indraprastha Institute of Information Technology, Delhi, Okhla Industrial Estate, Phase III, (Near Govind Puri Metro Station), New Delhi, India – 110020, Office: A-302 (R&D Block), Website: http://webs.iiitd.edu.in/raghava/. Mailing Address of Authors Sumeet Patiyal; Anjali Dhall; Gajendra P. S. Raghava.

## Abstract

DNA-protein interaction is one of the most crucial interactions in the biological system, which decide the fate of many processes such as transcription, regulation of gene expression, splicing, and many more. Though many computational approaches exist that can predict the DNA interacting residues from the protein sequences, there is still a significant opportunity for improvement in terms of performance and accessibility. In this study, we have downloaded the benchmark dataset from method hybridNAP and recently published method ProNA2020, for training and validation purposes, that comprise 864 and 308 proteins, respectively. We have implemented CD-HIT software to handle the redundancy with 30% identity, and left with 646 proteins for training and 46 proteins for validation purposes, in which the validation dataset do not share more than 30% of sequence identity with the training dataset. We have generated amino acid binary profiles, physicochemical-properties based binary profiles, PSSM profiles, and a combination of all profiles described as hybrid feature. 1D-CNN based model performed best as compared to other models for each set of features. The model developed using amino acid binary profile achieved AUROC of 0.83 and 0.74 for training and validation dataset. Using physicochemical properties based binary profile, model attained AUROC of 0.86 and 0.73 for training and validation dataset. Model generated using PSSM profile resulted in the better performance with AUROC 0.91 and 0.74 for training and validation dataset. And, model developed using hybrid of all features performed best with AUROC of 0.91, and 0.79 for training and validation dataset, respectively. We have compared our method’s performance with the current approach and shown improvements. We have included the best-performing models in the standalone and web server accessible at https://webs.iiitd.edu.in/raghava/dbpred. DBPred is an effective approach to predict the DNA interacting residues in the protein using its primary structure.

## Introduction

In every living organism, life is entirely dependent on a particular type of molecular interactions, such as DNA-protein, RNA-protein, protein-protein interactions, etc. These interactions perform several biological functions in the cells of the living organisms [1]. The DNA-protein interactions are the fundamental type of interactions for almost all biological activities and processes, such as transcription, gene expression regulation, repair, packaging of chromosomal DNA, and splicing [2-5]. Several experimental methods are used to confirm the interactions between protein and DNA-binding residues. The availability of experimental data on 3D structures of protein-DNA complexes and binding residues; supports biologists and researchers to reveal the essential knowledge on protein-DNA interactions such as conformational changes of DNA molecules, the importance of hydrogen bonds, amino-acid properties, electrostatic, van der Waals interactions, etc [6-15].

Due to the advancements of high-throughput sequencing a huge amount of experimentally curated DNA-proteins interaction data have registered in the protein data bank (PDB) [16]. But, identification of DNA-binding residues from the empirical data is very challenging, time-consuming, and costly process. Therefore, from the last few decades, several attempts have been made for the prediction of DNA-binding residues using computational methods [2, 17-19]. These tools are majorly divided into four categories, i.e., sequence-based methods [20], structure-based methods [21, 22], evolutionary methods [23] and hybrid methods which used both structure and sequence information [24, 25]. Several machine learning based methods like, BindN (PDNA-62 dataset) [18], BindN+ (PDNA-62 dataset) [26], BindN-RF (PDNA-62 dataset) [27], MetaDBSite (PDNA-316 dataset), DP-Bind (62 protein-DNA complexes) [20] have been developed for the identification of DNA-binding residues. But, the major limitation of these methods is that they build on a very small dataset of protein-DNA complexes. Whereas, some most popular methods, such as HybridNAP [28] (19987 DNA-binding residues), DRNApred (8791 DNA-binding residues) [29], ProNA2020 [30] which uses huge amount of data to develop the prediction models.

In the current study, we use this dataset in order to generate better prediction models for the identification of DNA-binding/non-binding residues. We introduce a new method named “DBpred” which uses deep learning and machine learning approaches for the accurate prediction of DNA-binding residues in the protein sequence. The benchmark dataset is consist of 864 annotated protein sequences (i.e., 16511 interaction and 307581 non-interacting residues) taken from hybridNAP and ProNA2020 benchmark dataset. This method uses amino-acid binary profiles, physicochemical binary profiles and evolutionary information for the development of prediction models. The machine learning models implemented on various classifiers, such as Random Forest (RF), Decision Tree (DT), eXtreme Gradient Boosting (XGB), Logistic Regression (LR), Gaussian Naive Bayes (GNB). Further, we have used deep learning approach (ID-CNN) for the precise prediction of DNA binding residues using primary sequence information. To serve the scientific community working in this era, we provide freely available webserver at https://webs.iiitd.edu.in/raghava/dbpred/, and standalone package at https://webs.iiitd.edu.in/raghava/dbpred/stand.php.

## Materials and Methods

### Dataset Creation

We have downloaded the dataset from the hybridNAP webserver [28] and recently published article ProNA2020 [30], which consists of 864 and 308 annotated protein sequences, respectively. Then, we have utilized the CD-HIT software [31] on these datasets to handle the redundancy with the standards of 30% sequence identity, and obtained 646 sequences in the training dataset, and 46 in the validation dataset, where the sequences in the validation dataset do not share more than 30% similarity with the sequences in the training dataset. Finally, we left with 15636 DNA-interacting and 298503 non-interacting residues in the training dataset, and 965 interacting and 9911 non-interacting residues in the validation dataset.

### Pattern Size

The overlapping patterns for each sequence with length 17 are generated using in-house python scripts. The central or 9th residue is taken as the representative of the obtained patterns. The pattern is specified as a positive segment if the central residue is DNA-interacting, else non-interacting or negative segment. In order to handle the terminal residues, eight counterfeit residues using the formula (N-1)/2 (where N represents the pattern length which is 17), as “X” are added at both sides of the protein sequences, as shown in Figure 1 along with the complete workflow for this study.

**Figure 1:**
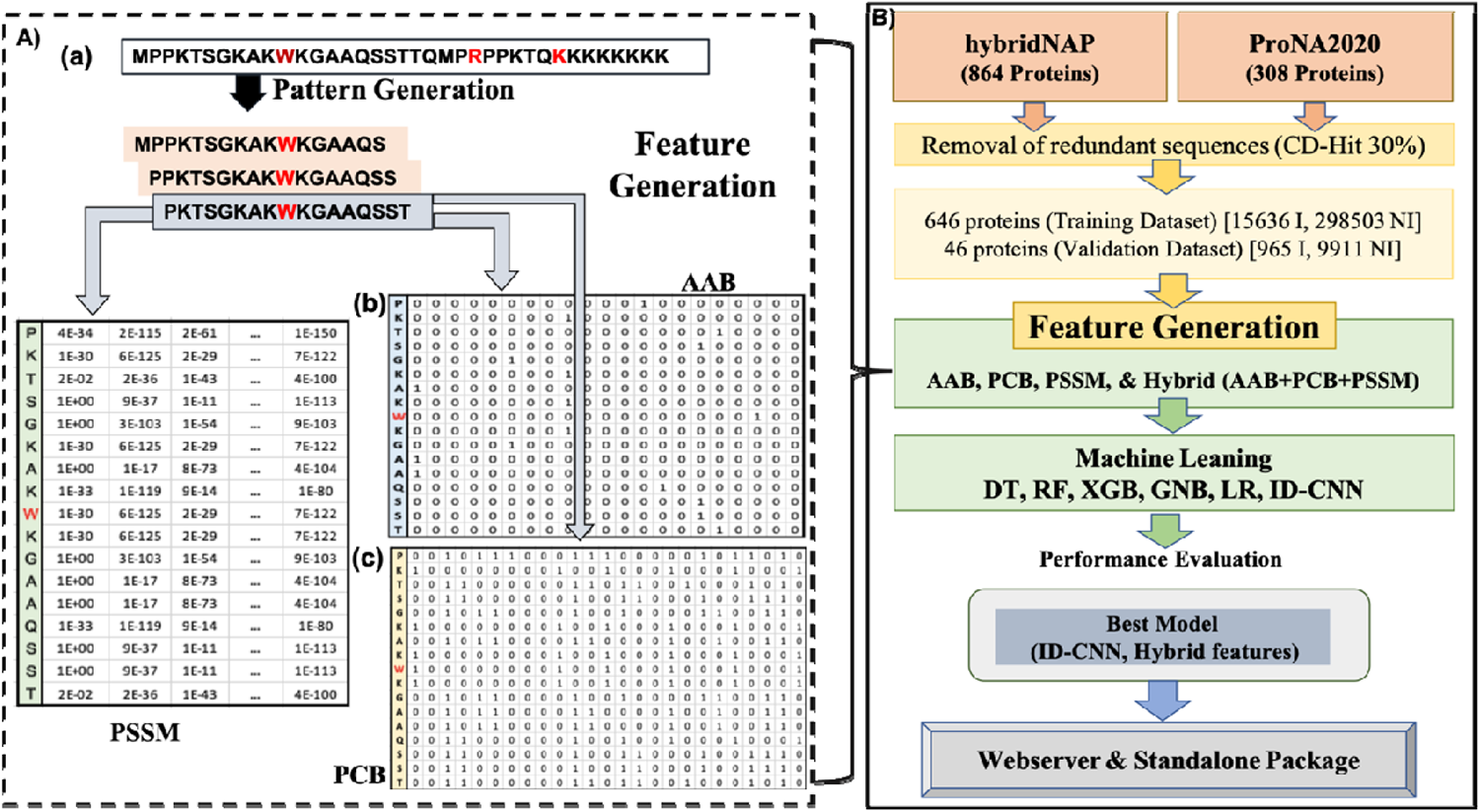
Comprehensive workflow and feature generation. A) Feature generation for patterns of length 17, DNA-interacting residues are shown in red-colour (e.g. W,R,K), positive pattern is shown in grey colour with ‘W’ as the central residue flanked by eight residues in each side, and the respective overlapping negative patterns are shown in peach colour. a) Generation of overlapping patterns of length 17, b) generation of amino acid binary profile (AAB) for each pattern, c) generation of physicochemical based binary profile (PCB), d) generation of PSSM profile for each pattern. B) Comprehensive workflow for DBPred.

### Percent Composition

In order to explore the nature of amino acid residues involved in the interaction with DNA, we have calculated the amino acid composition, residue propensity, and physicochemical properties-based composition. The percent amino acid composition was calculated using equation 1, which tells the abundance of residues in interaction. The residues propensity is computed using equation 2, which indicates the preference or non-preference of the particular type of residues in the DNA binding site. The functionality of residues is based on their sole physicochemical properties, and hence we have determined physicochemical properties-based composition for 25 distinct properties using equation 3. The properties that we have considered are positively charged, negatively charged, neutrally charged, polar, non-polar, aliphatic, cyclic, aromatic, acidic, basic, hydrophobic, hydrophilic, hydroxylic, sulphur-content, helix, strands, coil, buried, exposed, intermediate solvent accessibility, tiny, small, and large. All the percent compositions were calculated using the Pfeature package [32].

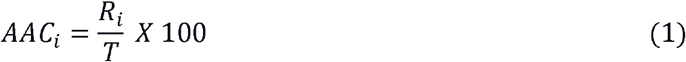

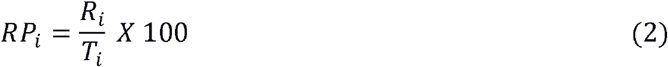

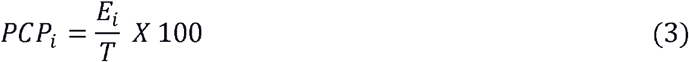

where, AAC_i_, RP_i_, and PCP_i_ is amino-acid composition, propensity score of residue i, respectively; PCP_i_ is composition of physicochemical property of type i;

R_i_ denotes the number of residues of type i;

T denotes the total number of residues;

T_i_ denotes the total number of residues of type i (DNA-interacting and non-interacting);

E_i_ denotes number of residues possessing physicochemical property of type i.

### Binary Profile

We have calculated two varieties of binary profiles for each pattern, such as amino acid binary profile (AAB) and physicochemical properties based binary profile (PCB). The features were calculated by modifying Pfeature [32] scripts. In AAB, each amino acid is represented by the vector size of length 21, for instance, A is described as 1,0,0,0,0,0,0,0,0,0,0,0,0,0,0,0,0,0,0,0,0; which comprises of 20 natural amino acids and one dummy variable, whereas X is denoted as 0,0,0,0,0,0,0,0,0,0,0,0,0,0,0,0,0,0,0,0,0 [33]. Therefore, each pattern is represented by the vector size of 357 (17*21). In PCB, each amino acid is designated by the vector of size 25; for instance, A is denoted by 0,0,1,0,1,1,0,0,0,0,1,1,0,0,0,0,1,0,0,1,0,0,1,1,0; where each position exhibits a particular physicochemical property, and each element denotes the presence (1) or absence (0) of that property. Therefore, the resulting vector for each property is of length 425 (17*25), whereas for X, all the elements are 0.

### PSSM Profile

The third feature that we have used is the evolutionary or <u>P</u>osition-<u>S</u>pecific <u>S</u>coring <u>M</u>atrix (PSSM) profile [34]. The PSSM profile was generated employing PSI-BLAST [35] by using the SwissProt database [36], against which each sequence is searched. The Parameters used for running PSI-BLAST were three iterations, with e-value as 1e-3. Further, the profile was normalized using equation 4. The final matrix for each sequence is of dimension Nx21, where N is the length of the protein sequence, and each pattern is depicted as the vector of length 357 (17*21).

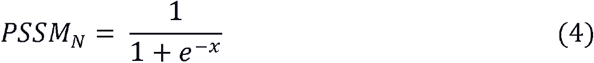

where, PSSM_N_ normalized value, x is the PSSM score.

### Machine Learning Predictors

We have implemented the python library scikit-learn based traditional machine learning and a one-dimensional CNN-based classifier using the TensorFlow library to develop the prediction models. In the conventional approach, we have implemented various classifiers, such as Random Forest (RF), Decision Tree (DT), eXtreme Gradient Boosting (XGB), Logistic Regression (LR), and Gaussian Naive Bayes (GNB), to develop the prediction model.

### Five-Fold Cross-Validation and Performance Evaluation

In order to avoid overfitting, biasness and to evaluate the performance of the generated prediction models, we have implemented the five-fold cross-validation. In this method, all the dataset is divided into five non-overlapping sets, four out of five sets are used for the training purpose, and the fifth set is kept for testing. The same process is repeated five times so that each set gets the chance to be used as the testing dataset only once. The overall performance would be the mean of the performances of five iterations [37-39].

In this study, we have calculated various threshold-dependent and threshold-independent parameters in order to evaluate prediction models. Threshold -dependent parameters include sensitivity (sens, equation 5), which signify the percentage of correctly predicted DNA-interacting residues; specificity (spec, equation 6) explains the proportion of correctly predicted DNA non-interacting residues; accuracy (acc, equation 7) defines the percentage of correctly predicted DNA-interacting and non-interacting residues; and Matthews Correlation Coefficient (MCC, equation 8), which exhibits the correlation between observed and predicted values. On the other hand, the threshold-independent parameter includes Area Under Receiver Operating Characteristics (AUROC), which is the plot between True Positive Rate (TPR) and False Positive Rate (FPR). The module of the R named “pROC” was used to plot the AUROC curve [40]. The equations for threshold dependent parameters are as follows:

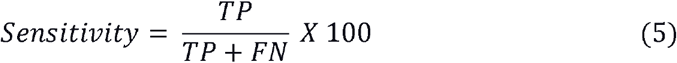

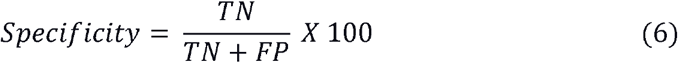

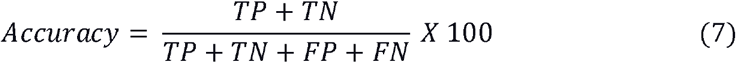

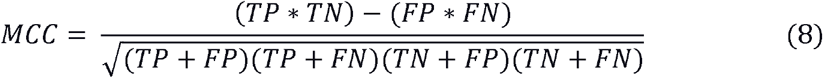

Where FP, FN, TP, and TN are false positive, false negative, true positive, and true negative, respectively.

## Results

### Compositional Analysis

We have performed the compositional analysis to understand the interactions of the DNA to the protein residues. We have analyzed the amino acid composition of DNA-interacting and non-interacting residues in DNA interacting proteins. As shown in Figure 2, DNA-interacting residues are rich in H, K, N, R, and Y, whereas, A, D, I, L, and P are abundant among non-interacting residues. In order to explore the preference of residues in the DNA-binding site, we have calculated the propensity of each residue, which exhibits that K, R, W, and Y are most favoured in the DNA-binding site, as shown in Figure 3. We have also analyzed the residues’ properties involved in interaction with DNA and found that positively charged, basic, hydrophilic, possessing helix secondary structure, and large are more abundant in DNA-interacting residues shown in Figure 4.

**Figure 2:**
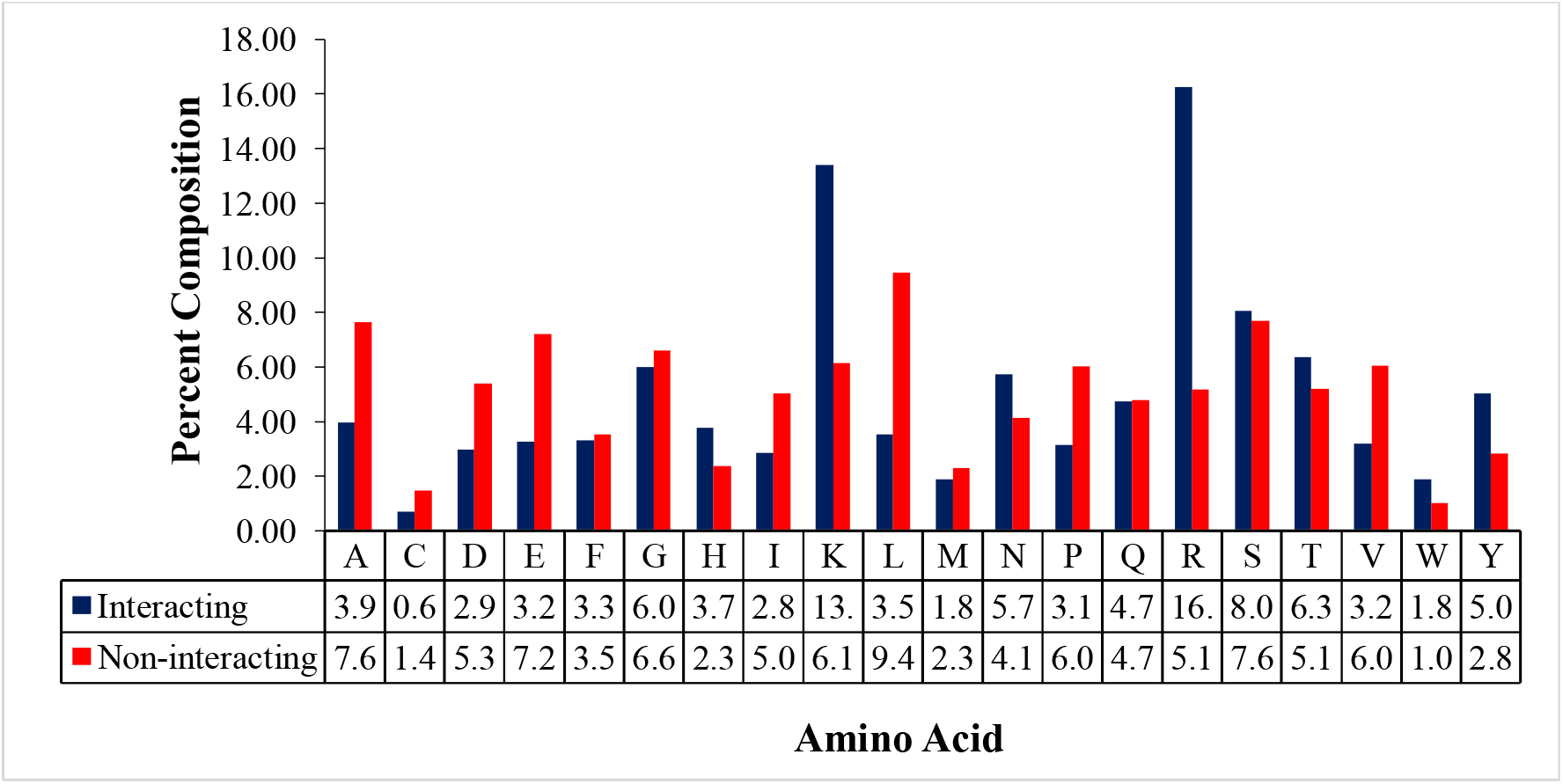
Percent composition of DNA-interacting and non-interacting residues

**Figure 3:**
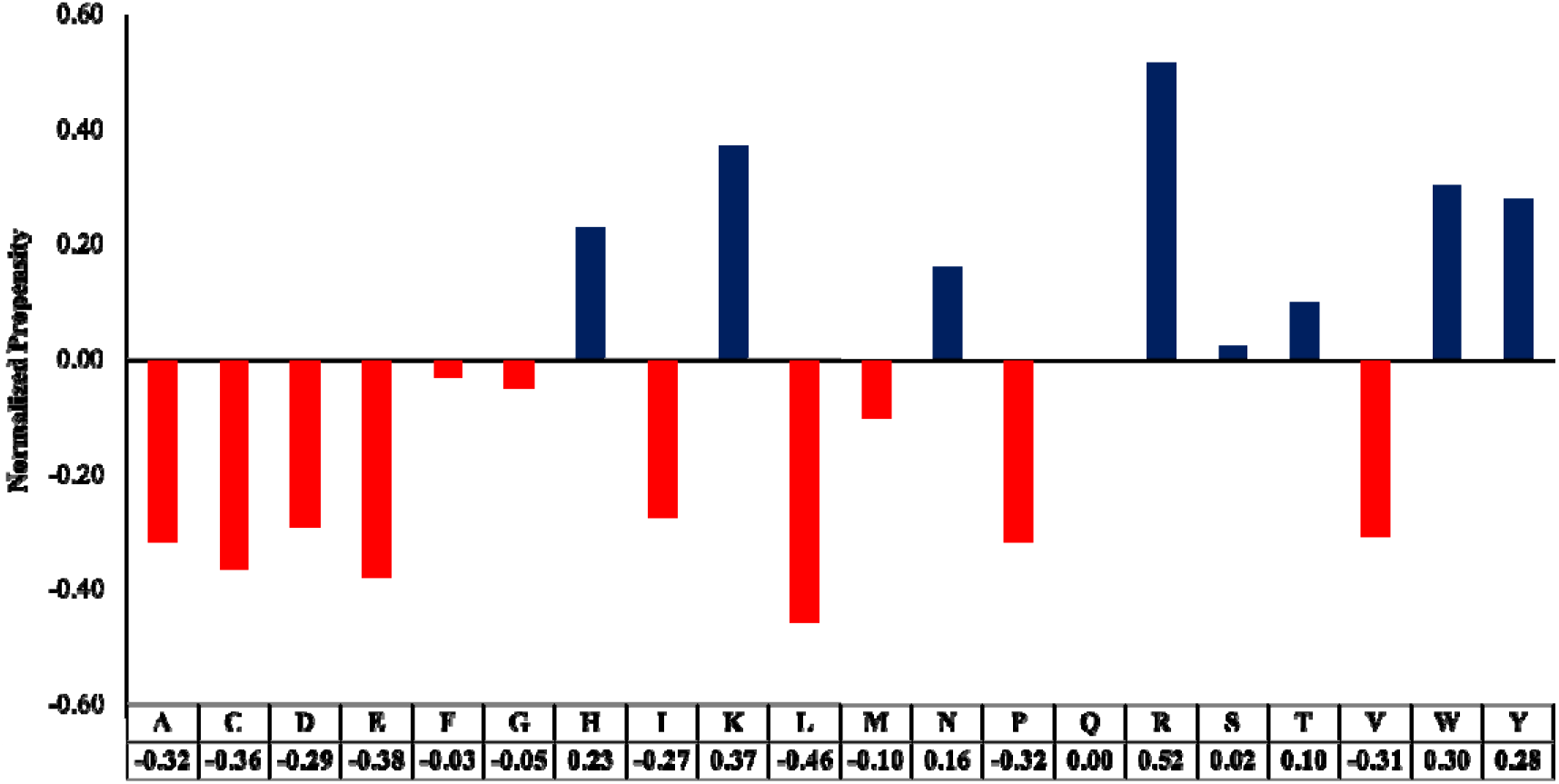
Normalized propensity scores DNA-interacting and non-interacting residues

**Figure 4:**
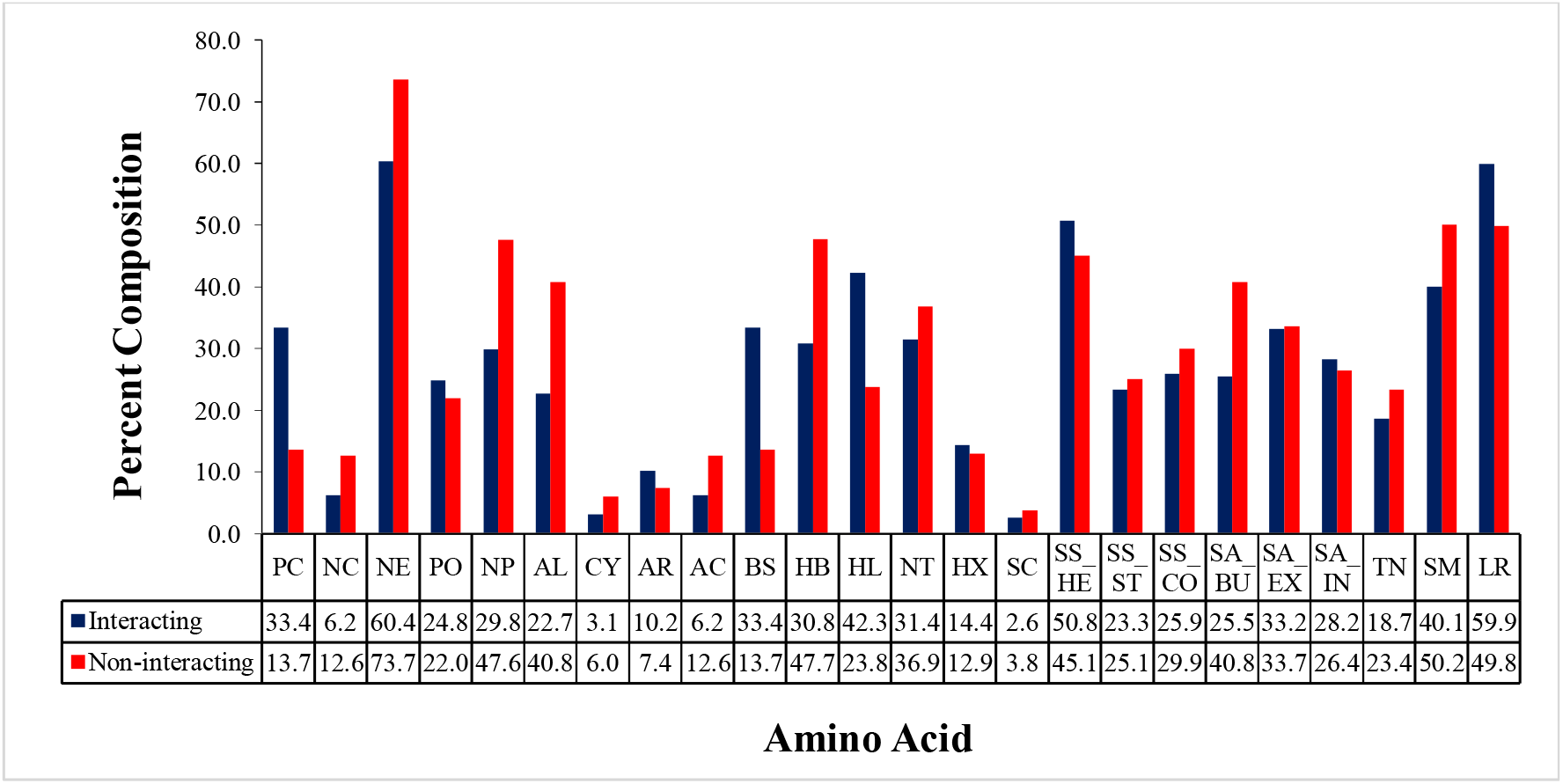
Percent composition of physicochemical properties in DNA-interacting and non-interacting residues

### Performance based on Amino Acid Binary Profile

In order to develop the prediction models, we have generated amino acid binary profile, as it captures the compositional as well as positional information of each residue. We have generated the binary profile for the training dataset consisting of 15636 patterns for DNA-interacting and 298503 non-interacting patterns; and the validation dataset comprises 965 DNA-interacting and 9911 non-interacting patterns. The best result for each classifier is shown in table 1. As shown in Table 1, the one-dimensional CNN-based classifier performed best among all the other classifiers with AUROC 0.83 and MCC 0.25 for training dataset, and AUROC 0.74 and MCC 0.21 for the validation dataset.

**Table 1:**
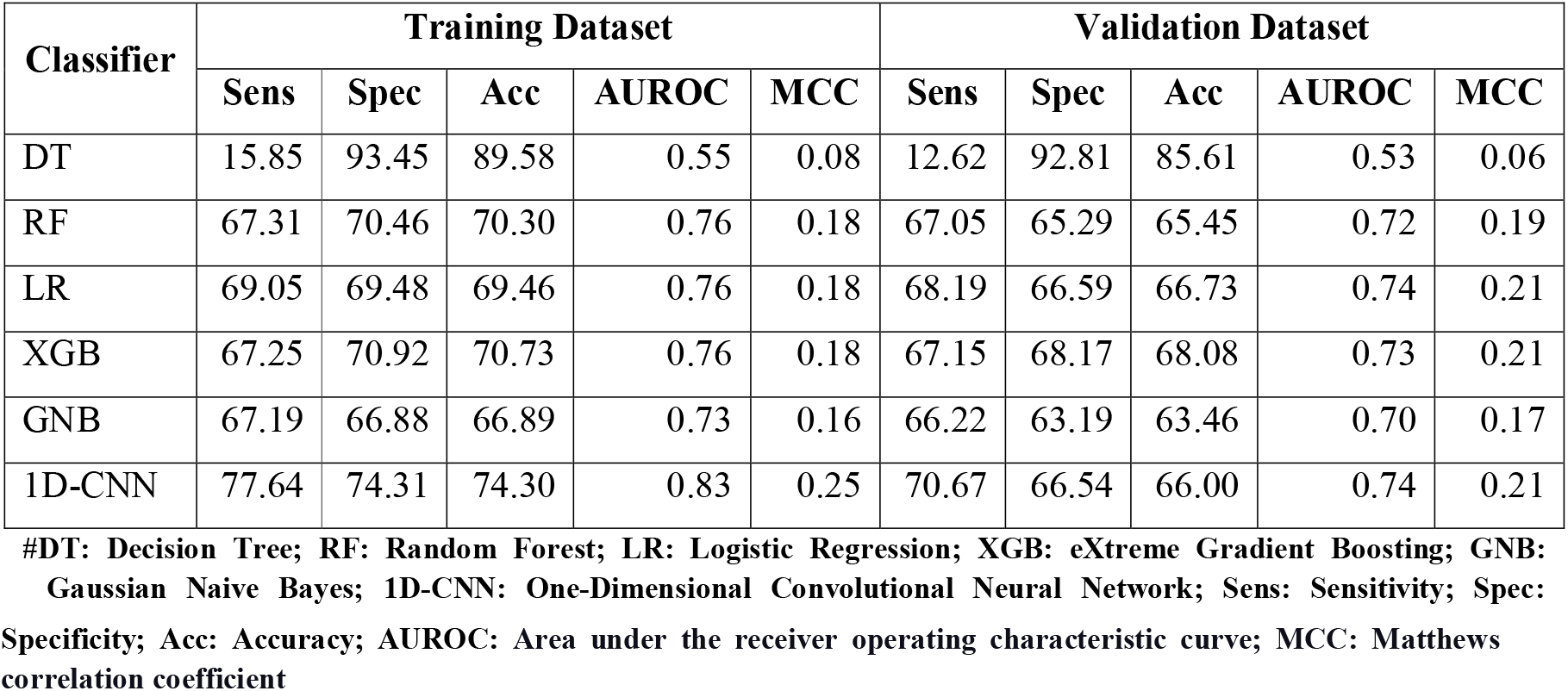
Performance of various classifiers using amino acid binary profile.

| Classifier | Training Dataset |  |  |  |  | Validation Dataset |  |  |  |  |
| --- | --- | --- | --- | --- | --- | --- | --- | --- | --- | --- |
|  | Sens | Spec | Acc | AUROC | MCC | Sens | Spec | Acc | AUROC | MCC |
| DT | 15.85 | 93.45 | 89.58 | 0.55 | 0.08 | 12.62 | 92.81 | 85.61 | 0.53 | 0.06 |
| RF | 67.31 | 70.46 | 70.30 | 0.76 | 0.18 | 67.05 | 65.29 | 65.45 | 0.72 | 0.19 |
| LR | 69.05 | 69.48 | 69.46 | 0.76 | 0.18 | 68.19 | 66.59 | 66.73 | 0.74 | 0.21 |
| XGB | 67.25 | 70.92 | 70.73 | 0.76 | 0.18 | 67.15 | 68.17 | 68.08 | 0.73 | 0.21 |
| GNB | 67.19 | 66.88 | 66.89 | 0.73 | 0.16 | 66.22 | 63.19 | 63.46 | 0.70 | 0.17 |
| 1D-CNN | 77.64 | 74.31 | 74.30 | 0.83 | 0.25 | 70.67 | 66.54 | 66.00 | 0.74 | 0.21 |

#DT: Decision Tree; RF: Random Forest; LR: Logistic Regression; XGB: eXtreme Gradient Boosting; GNB: Gaussian Naive Bayes; 1D-CNN: One-Dimensional Convolutional Neural Network; Sens: Sensitivity; Spec:

### Performance based on Physicochemical Properties Binary Profile

We have also used the binary profiles based on physicochemical properties for the first time in the literature to develop the prediction models. As shown in Table 2, 1D-CNN based model has outperformed all the other classifiers with AUROC 0.86 and MCC 0.27 for the training dataset, and AUROC 0.73 and MCC 0.20 on the validation dataset.

**Table 2:**
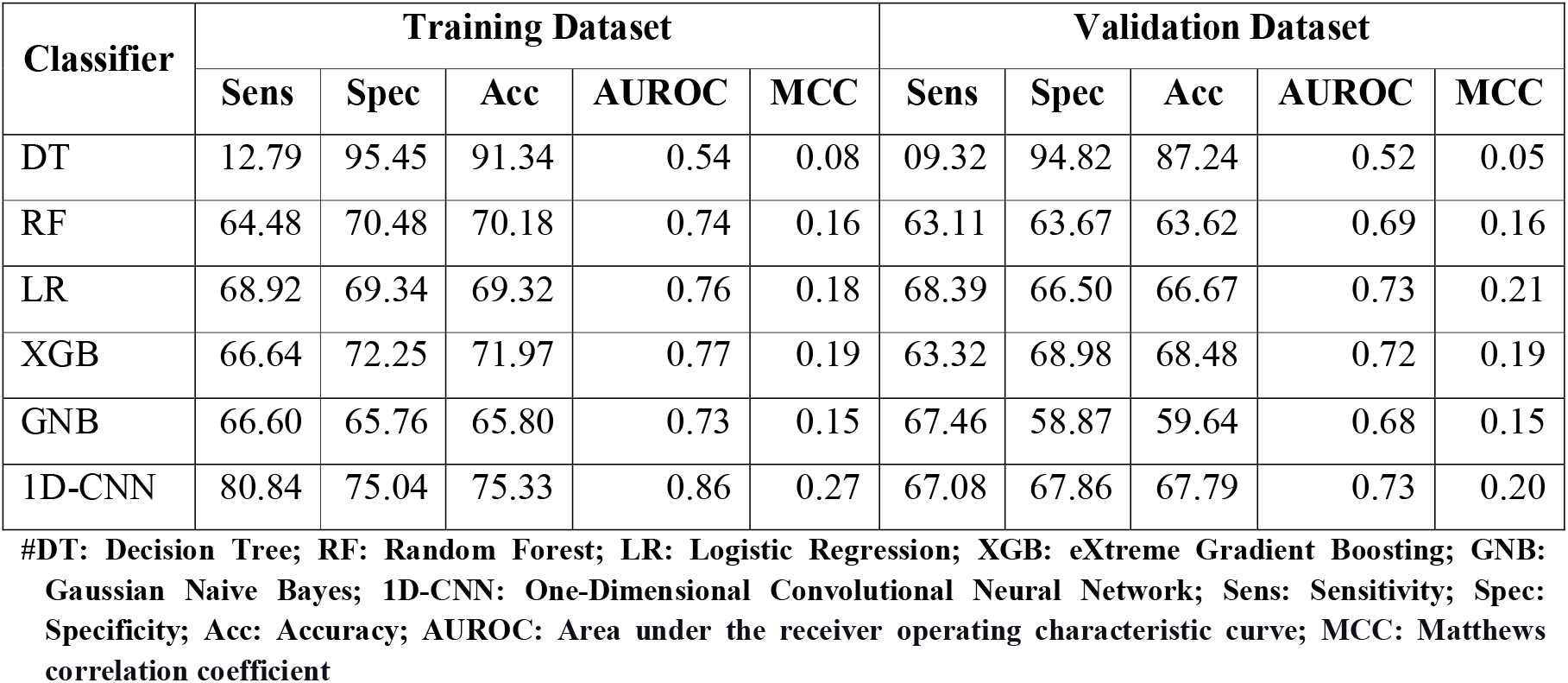
Performance of various classifiers using physicochemical properties based binary profile.

| Classifier | Training Dataset |  |  |  |  | Validation Dataset |  |  |  |  |
| --- | --- | --- | --- | --- | --- | --- | --- | --- | --- | --- |
|  | Sens | Spec | Acc | AUROC | MCC | Sens | Spec | Acc | AUROC | MCC |
| DT | 12.79 | 95.45 | 91.34 | 0.54 | 0.08 | 09.32 | 94.82 | 87.24 | 0.52 | 0.05 |
| RF | 64.48 | 70.48 | 70.18 | 0.74 | 0.16 | 63.11 | 63.67 | 63.62 | 0.69 | 0.16 |
| LR | 68.92 | 69.34 | 69.32 | 0.76 | 0.18 | 68.39 | 66.50 | 66.67 | 0.73 | 0.21 |
| XGB | 66.64 | 72.25 | 71.97 | 0.77 | 0.19 | 63.32 | 68.98 | 68.48 | 0.72 | 0.19 |
| GNB | 66.60 | 65.76 | 65.80 | 0.73 | 0.15 | 67.46 | 58.87 | 59.64 | 0.68 | 0.15 |
| 1D-CNN | 80.84 | 75.04 | 75.33 | 0.86 | 0.27 | 67.08 | 67.86 | 67.79 | 0.73 | 0.20 |

#DT: Decision Tree; RF: Random Forest; LR: Logistic Regression; XGB: eXtreme Gradient Boosting; GNB: Gaussian Naive Bayes; 1D-CNN: One-Dimensional Convolutional Neural Network; Sens: Sensitivity; Spec: Specificity; Acc: Accuracy; AUROC: Area under the receiver operating characteristic curve; MCC: Matthews correlation coefficient

### Evolutionary information based performance

As shown by the literature in the past, evolutionary information captures more information than any other method. We have described the evolutionary information as the PSSM profile. We have developed various prediction models by using normalized PSSM profile as the input feature, and the performance of each classifier is exhibited in Table 3. 1D-CNN-based classifier exceeded other classifiers’ performance, 0.91 AUROC and 0.34 MCC for the training dataset, and AUROC of 0.74 and MCC of 0.21 for the validation dataset.

**Table 3:** Performance of various classifiers using PSSM profile.

| Classifier | Training Dataset |  |  |  |  | Validation Dataset |  |  |  |  |
| --- | --- | --- | --- | --- | --- | --- | --- | --- | --- | --- |
|  | Sens | Spec | Acc | AUROC | MCC | Sens | Spec | Acc | AUROC | MCC |
| DT | 20.31 | 95.67 | 91.92 | 0.58 | 0.16 | 13.26 | 94.47 | 87.27 | 0.54 | 0.09 |
| RF | 73.47 | 71.69 | 71.78 | 0.81 | 0.21 | 73.06 | 62.46 | 63.41 | 0.74 | 0.21 |
| LR | 72.92 | 71.94 | 71.99 | 0.80 | 0.21 | 69.33 | 67.98 | 68.10 | 0.75 | 0.22 |
| XGB | 76.78 | 73.96 | 74.10 | 0.84 | 0.24 | 72.12 | 67.51 | 67.92 | 0.77 | 0.24 |
| GNB | 66.68 | 63.56 | 63.71 | 0.70 | 0.14 | 64.87 | 56.24 | 57.01 | 0.63 | 0.12 |
| 1D-CNN | 88.09 | 79.45 | 79.88 | 0.91 | 0.34 | 64.89 | 69.87 | 69.43 | 0.74 | 0.21 |

#DT: Decision Tree; RF: Random Forest; LR: Logistic Regression; XGB: eXtreme Gradient Boosting; GNB: Gaussian Naive Bayes; 1D-CNN: One-Dimensional Convolutional Neural Network; Sens: Sensitivity; Spec: Specificity; Acc: Accuracy; AUROC: Area under the receiver operating characteristic curve; MCC: Matthews correlation coefficient

### Performance based on combined features

The combined features were generated by concatenating the amino acid binary profile, physicochemical properties-based binary profile, and PSSM profile in the column-wise manner for each pattern, which generated a vector of length 1175. A wide range of classifiers was used to develop the prediction method, and the 1D-CNN-based classifier performed best among all the classifiers with 0.91 AUROC and 0.37 MCC on the training dataset, and 0.79 AUROC and 0.32 MCC on the validation dataset, as shown in Table 4.

**Table 4:** Performance of various classifiers using combined features.

| Classifier | Training Dataset |  |  |  |  | Validation Dataset |  |  |  |  |
| --- | --- | --- | --- | --- | --- | --- | --- | --- | --- | --- |
|  | Sens | Spec | Acc | AUROC | MCC | Sens | Spec | Acc | AUROC | MCC |
| DT | 16.81 | 95.49 | 91.57 | 0.56 | 0.12 | 15.34 | 94.91 | 87.84 | 0.55 | 0.18 |
| RF | 74.27 | 72.04 | 72.15 | 0.81 | 0.22 | 70.98 | 63.02 | 63.73 | 0.75 | 0.20 |
| LR | 74.28 | 73.36 | 73.40 | 0.81 | 0.23 | 70.88 | 69.18 | 69.33 | 0.77 | 0.29 |
| XGB | 68.69 | 67.38 | 67.44 | 0.74 | 0.17 | 69.95 | 62.62 | 63.27 | 0.72 | 0.19 |
| GNB | 69.99 | 68.11 | 68.20 | 0.75 | 0.18 | 66.84 | 62.70 | 63.06 | 0.70 | 0.17 |
| 1D-CNN | 83.74 | 83.50 | 83.51 | 0.91 | 0.37 | 70.78 | 78.40 | 77.72 | 0.79 | 0.32 |

#DT: Decision Tree; RF: Random Forest; LR: Logistic Regression; XGB: eXtreme Gradient Boosting; GNB: Gaussian Naive Bayes; 1D-CNN: One-Dimensional Convolutional Neural Network; Sens: Sensitivity; Spec: Specificity; Acc: Accuracy; AUROC: Area under the receiver operating characteristic curve; MCC: Matthews correlation coefficient

### Comparison with the existing methods

In order to concede the newly developed method, its comparison with the existing methods is of uttermost importance. The comparison conveys the merits and demerits of the newly developed method. Since there are many existing methods for predicting DNA-binding residues in a protein [26-28,30], a comprehensive comparison is must to understand the benefits of the newly developed method “DBPred”. We have calculated Physico-chemical properties based binary profile as the new feature in DBPred, other than amino acid binary profile and evolutionary information, but its performance is equivalent to the performance using the amino acid binary profile. We have developed models on individual features and their combination; the model developed on the combined feature outperforms the models developed on the individual features mentioned in Tables 1, 2, 3, and 4. The performance of existing methods along with the datasets used is shown in table 5. The performances are reported in terms of AUROC and accuracy. In some of the methods, such as BindN-RF [27], DBindR [41], BindN+ [26], DNABR [42], and PDNASite [43], the performance is relatively higher, which could be due to the overfitting of the model, since the used datasets are smaller in size. On the other hand, recent methods like TargetDNA [44], HybridNAP [28], funDNApred [45], iProDNA-CapsNet [46], and ProNA2020 [30] have used larger datasets as compared to the previous methods; the DBPred had used the equivalent dataset to the recently developed methods and outperformed them. As shown in Table 5, most methods have provided the webserver facility, but many are non-functional now. Like most of the existing methods, DBPred has furnished the webserver service, which incorporates many facilities for the users, such as, they can provide multiple sequences at a time for the prediction, various modules have been provided based on the types of feature used to develop models. The website is designed using an HTML5 responsive template, which is compatible with all the latest devices, such as mobile, iPad, tablets, laptops, and desktops. This is the age of genomics where a user can wish to predict the DNA-binding residues in the whole proteome, which is quite heavy for the webserver to handle; hence we have developed the python-and Perl-based standalone that can be downloaded and run on the local machines irrespective of their operating systems and can be used in the absence of internet.

**Table 5:** Comparison of performance of various existing methods with DBPred.

| Method | Year | Dataset | Redundancy | AUROC | Accuracy | MCC | Webserver/<br>Standalone |
| --- | --- | --- | --- | --- | --- | --- | --- |
| DBS-Pred [47] | 2004 | 62 | 25% | - | 64.00% | - | W* |
| DBS-PSSM [48] | 2005 | 62 | 25% | - | 67.10% | - | W* |
| Pro-DNA [49] | 2005 | 115 | - | - | 79.00% | - | W* |
| BindN [18] | 2006 | 62 | 25% | 0.75 | 70.31% | - | W* |
| DP_Bind [50] | 2006 | 62 | 25% | 0.84 | 76.00% | 0.45 | W<sup>#</sup> |
| DNABindR [51] | 2006 | 171 | 30% | - | 78.00% | 0.28 | - |
| DP-Bind [20] | 2007 | 62 | 25% | - | 77.20% | - | W<sup>#</sup> |
| BindN-RF [27] | 2009 | 62 | 25% | 0.86 | 78.20% | - | W* |
| DBindR [41] | 2009 | 374 | 25% | 0.91 | 91.41% | 0.70 | W* |
| BindN+ [26] | 2010 | 62 | 25% | 0.86 | 79.00% | 0.44 | W* |
| MetaDBSite [52] | 2011 | 316 | 30% | - | 77.00% | 0.32 | W* |
| DNABR [42] | 2012 | 337 | 25% | 0.88 | 93.04% | 0.66 | W* |
| DNABind [24] | 2013 | 206 | 25% | 0.81 | - | 0.38 | W<sup>#</sup> |
| SPOT-Seq (DNA) [53] | 2014 | 179 | 35% | - | 88.00% | 0.52 | W<sup>#</sup> |
| Wang et. al [54] | 2014 | 272 | 30% | 0.86 | 85.40% | 0.72 | - |
| PDNASite [43] | 2016 | 286 | 25% | 0.93 | 85.11% | 0.58 | W* |
| TargetDNA [44] | 2017 | 543 | 30% | 0.82 | 76.42% | 0.30 | W<sup>#</sup> |
| HybridNAP [28] | 2017 | 864 | - | 0.69 | - | 0.19 | W<sup>#</sup> |
| funDNApred [45] | 2018 | 864 | - | 0.72 | 84.50% | 0.19 | W<sup>#</sup> |
| iProDNA-CapsNet [46] | 2019 | 543 | 30% | 0.83 | 76.02% | 0.29 | W<sup>#</sup> |
| ProNA2020 [30] | 2020 | 308 | 20% | - | 81.00% | 0.42 | W<sup>#</sup>/S |
| DBPred | 2021 | 692 | 40% | 0.91 | 83.51% | 0.37 | W<sup>#</sup>/S |

### Web server implementation

In order to serve the scientific community, we have developed and executed our best models in the webserver “DBPred”, to predict the DNA-interacting residues in a protein using its primary structure information. The facilities provided by the webserver are available in various modules such as “Sequence,” “PSSM profile,” “Hybrid,” and “Standalone.” The description of each module is as follows.

### Sequence Module

In this module, the user is allowed to predict the DNA interacting residues in the query protein sequences by providing them in the FASTA format. This module enables users to choose the type of input features such as amino acid binary profile (AAB) and physicochemical properties based binary profile (PCB), with the desired threshold value vary between 0 and 1. The sequence module provides the facility to either paste or upload sequence(s) in the FASTA format. On the result page, DNA-interacting residue(s) in the protein sequence(s) are shown in red, whereas non-interacting residues are shown in black. The results are downloadable in different formats.

### PSSM Module

The PSSM module produces a PSSM profile for the query proteins, which is used as the input feature to predict the DNA-interacting potential of the residues in the submitted protein sequences. This module also authorizes the users to vary the probability threshold. The PSSM module permits the users to either paste the sequence(s) in the provided area or upload the file of sequences in the FASTA format. The result page exhibits the query protein sequences, where DNA-interacting residues are shown in red colour, and it also provides the facility to download the result in either txt, png, or pdf format.

### Hybrid Module

This module implements the hybrid of the features mentioned in the above modules, such as AAB, PCB, and PSSM, to predict the DNA-interacting residues in the query protein sequence(s). This module also provides the facilities granted by the above modules, such as selecting the desired probability threshold, single or multiple sequences at a time, and paste or upload file alternatives. The DNA-interacting residues would be shown in red colour with bigger font in the protein sequence(s), on the result page, with the option of downloading the results in either pdf, txt, or png format.

### Standalone

Other than the web server, we have also developed the Perl-and python-based standalone, which can be used in offline mode. These standalone versions have the best models executed in the back-end. The standalone takes the sequence(s) in the FASTA format as the input and provides the annotated file as output. This module permits the users to download the standalone versions, and it also provides the stepwise execution of the standalone in the docker.

## Discussion

Methods with the ability to identify the DNA-interacting sites on protein can broadly be classified into one of the three classes such as sequence-based, structure-based, and hybrid approaches [23, 55]. The limitation of the structure-based or hybrid methods is their dependency on the protein structural information, which limits their application, as determination of the protein structure is a costly, time-consuming, and very complex process [33]. On the other hand, sequence information in various databases is growing exponentially, enhancing the application of sequence-based methods with reliable performance. In this study, we have made a systematic attempt to develop a prediction method that can predict the DNA-interacting residues in a protein sequence. We have explored various properties of DNA-interacting residues such as amino acid composition, physicochemical properties composition, propensities of the residues and developed different prediction models using multiple machine learning classifiers such as DT, RF, XGB, LR, GNB, and 1D-CNN. 1D-CNN-based method using a combination of amino acid binary profile, physicochemical properties based binary profile, and PSSM profile as input features performed best among the other classifiers. To the best of our knowledge, our approach has exceeded the performance of the existing methods, such as hybridNAP has reported AUROC 0.69 whereas our method is exhibiting AUROC of 0.91; similarly recently published method ProNA2020 has shown the accuracy of 81% while DBPred is showing the accuracy of 83.51%. We believe that DBPred can be an efficient tool for correctly predicting DNA-interacting residues in a protein sequence. To serve the scientific community, we have developed the standalone and web server “DBPred” to assist biologists in the finding of DNA-interacting residues for the sake of annotation and functional analysis. DBPred is freely available and accessible on https://webs.iiitd.edu.in/raghava/dbpred/.

## Conflict of inter est

The authors declare no competing financial and non-financial interests.

## Authors’ contributions

SP and GPSR collected and processed the datasets. SP and GPSR implemented the algorithms and developed the prediction models. SP, AD, and GPSR analysed the results. SP created the back-end of the web server and AD created the front-end user interface. SP, AD, and GPSR penned the manuscript. GPSR conceived and coordinated the project. All authors have read and approved the final manuscript.

## Acknowledgements

Authors are thankful to the Department of Bio-Technology (DBT) and Department of Science and Technology (DST-INSPIRE) for fellowships and the financial support and Department of Computational Biology, IIITD New Delhi for infrastructure and facilities.

